# Realized flower constancy in bumble bees: optimal foraging strategy balancing cognitive and travel costs and its possible consequences for floral diversity

**DOI:** 10.1101/2024.04.08.588636

**Authors:** Kentaro Takagi, Kazuharu Ohashi

## Abstract

1. Pollinating insects often exhibit flower constancy, i.e., the tendency to make consecutive visits to the same flower species while disregarding others. This behaviour is commonly attributed to the cost of retrieving visual or motor memories from long-term storage while switching between flowers with distinct colours and shapes. Accordingly, researchers often predict co-flowering species to exhibit significantly greater phenotypic diversity than random expectation, thereby minimizing heterospecific pollen transfer. However, field observations have not consistently supported this notion.
2. The observed inconsistencies may arise from variations in travel costs, which depend on the interaction between the forager’s constancy level and the spatial mixing of plant species. If species are evenly mixed, constant pollinators incur higher levels of travel cost due to frequent skipping of neighbouring flowers. In contrast, if species are patchily distributed, constant pollinators experience lower levels of travel cost, as most neighbours are of the same species. Considering this, ‘realized flower constancy’ may be determined as an optimal strategy for balancing cognitive and travel costs, which dynamically vary across different degrees of spatial species mixing. Here we test this possibility in indoor experiments with bumble bees foraging from two differently coloured artificial flowers (’species’) arranged at three mixing levels.
3. First, bees dramatically reduced flower constancy as species mixing increased, irrespective of flower spacing. Second, bees were less inclined to switch species after accumulating consecutive visits to one species, suggesting a rapid decay of another species’ information in short-term memory back to long-term storage. This effect may have additionally contributed to the increased flower constancy observed in species with patchy distributions. Third, bees showed minimal constancy for similarly coloured, evenly mixed flower species, suggesting that these flowers were operated with shared short-term memory. The constancy level was hardly affected by colour similarity when species were patchily distributed.
4. Results support our initial hypothesis that realized flower constancy reflects an optimal foraging strategy rather than a fixed outcome of cognitive limitation. Notably, bees’ constancy increased significantly with greater colour difference only when species were evenly mixed, suggesting a novel perspective: spatial mixing promotes the evolution and maintenance of floral diversity.

## 1 INTRODUCTION

Pollinating insects often exhibit flower constancy, i.e., the tendency to make consecutive visits to the same type of flowers even when other floral resources are available nearby (Chittka et al. 1999). Such persistent behaviour of pollinators confers clear advantages to animal-pollinated plants by preventing the reduction of male or female reproductive success through heterospecific pollen transfer (Morales & Traveset 2008; Moreira-Hernández & Muchhala 2019). In this light, it is crucial to elucidate the principles by which pollinators determine the degree of flower constancy in order to deepen our understanding of how floral diversity has evolved and been maintained through interactions with animals (Chittka et al. 2001; Jones 2001).

The most commonly accepted explanation for flower constancy is cognitive limitation, wherein pollinators must retrieve visual or motor memories from long-term storage each time they switch between flowers with distinct colours and shapes (Lewis 1986; Wilson & Stine 1996; Chittka et al. 1997, 1999; Goulson 2000; Ishii 2005; Raine & Chittka 2007; Ishii & Masuda 2014; Ishii & Kadoya 2016). Due to the time cost associated with this memory retrieval, pollinators tend to prefer the same type of flowers as those they have recently visited. Studies on bees, like those on vertebrates, have identified distinct temporal forms of memory, termed short-term and long-term memory (Menzel 1999; Greggers & Menzel 1993; Chittka et al. 1999). Long-term memory (LTM) stores substantial information for hours or more. However, before this information can be utilized, it must first be transferred to short-term memory (STM) or working memory (Chittka et al. 1999). Considering the limited capacity and duration of STM, this implies that bees will need to retrieve the same information from LTM each time they switch between flower types. For this reason, pollinators are believed to save time by temporarily specializing in a single type, thus avoiding the cognitive cost associated with alternating between flowers with distinct characteristics. In support of this notion, pollinators tend to exhibit greater constancy as differences in floral traits, especially colour, increase (Wilson & Stine 1996; Chittka et al. 2001; Gegear & Laverty 2005; de Jager et al. 2011; Hopkins & Rausher 2012).

Building on this concept, researchers often predict that flower constancy of pollinators will enhance floral phenotypic diversity in co-flowering communities, primarily through two processes: (1) ecological sorting, where flower constancy acts as a filter, shaping community assemblages of floral phenotypes by facilitating the establishment and persistence of species with distinct flowers from their surroundings (Gumbert et al. 1999), or (2) character displacement, where flower constancy exerts divergent selection pressures driving co-flowering species to evolve distinct floral phenotypes (Gumbert et al. 1999; Hopkins & Rausher 2012). Contrary to this prediction, however, previous investigations on floral colour composition in co-flowering communities have yielded varying outcomes, ranging from significant diversities to high similarities compared to what would be expected by chance, as well as instances of complete randomness (Gumbert et al. 1999; de Jager et al. 2011; Makino & Yokoyama 2015; Shrestha et al. 2019).

These inconsistencies in observation indicate that the level of flower constancy realized in field situations is not merely a fixed outcome of cognitive limitation but rather a conditional behaviour that responds to changes in some factor in the environment. While most studies on flower constancy have heavily focused on the effects of pollinators’ limited ability to switch between distinct memories within a short timeframe, it is often overlooked that flower constancy could be an adaptive behaviour in its own right, quickly adjustable based on the energetic value of different flower types (Grüter & Ratnieks 2011). In this study, we explore such potential adaptiveness of flower constancy within the framework of optimal foraging theory (Pyke 1984). To this end, we avoid directly inheriting the conventional term ‘flower constancy’, which is primarily attributed to the cost of memory retrieval, and instead introduce the new term ‘realized flower constancy’. We particularly focus on how travel costs for pollinators vary with the spatial mixing of plant species, from highly mixed to patchily distributed, and how their effects manifest in the realized flower constancy. Given that pollinators typically select the nearest neighbours when moving between plants and flowers (Pyke 1978; Ohashi et al. 2007), it seems reasonable to anticipate travel costs to be a major counteracting force against cognitive shortcuts.

The travel costs experienced by pollinators will depend on the interaction between their constancy level and the spatial mixing of plant species. If different species are evenly mixed, constant pollinators that circumvent the memory retrieval process will incur higher levels of travel costs due to frequent skipping of heterospecific neighbours. In such a trade-off situation, realized flower constancy will represent a compromise between cognitive and travel costs. In other words, realized flower constancy may drop to a level indistinguishable from nearest-neighbour movements in the absence of memory retrieval cost. If species are patchily distributed, on the other hand, constant pollinators will experience lower levels of travel costs, as neighbours are often conspecific. In this situation, pollinators will minimize both cognitive and travel costs simply by moving between the nearest neighbours, which results in high levels of realized flower constancy regardless of the presence of retrieval cost. Furthermore, the long consecutive visits to one species in patchy distributions may provide feedback to reinforce realized flower constancy toward this particular species. This is because persistent visits to one type of flower are likely to cause the information of another type to fade from STM to LTM (Wilson & Stine 1996; Chittka et al. 1999; Ishii 2006; Raine & Chittka 2007).

Here we conducted an indoor experiment with bumble bees to investigate the aforementioned possibilities. Specifically, we examined how bees change their levels of flower constancy in response to the varying degrees of mixing between two differently coloured artificial flowers. Our goal is to reframe current understandings of flower constancy from the viewpoint of optimal foraging theory, provide a conceptual framework for comprehending the variation in realized flower constancy across environments, and finally, discuss its implications for floral ecology and evolution. In this study, we addressed the following three questions: (i) How does realized flower constancy change with increasing species mixing and spacing between flowers? (ii) Do long consecutive visits to the same species in patchy distributions increase the retrieval cost? (iii) How does the spatial mixing of species affect the impact of floral colour differences on realized flower constancy?

## 2 MATERIALS AND METHODS

### 2.1 Bees and flight cages

We worked indoors in either of two flight cages, measuring 400 × 400 × 200 (height) cm and 250 × 200 × 200 (height) cm, respectively. Temperature ranged from 21 to 25℃. Both cages were made of steel pipes and greenhouse vinyl, and their floors were covered with green carpet. Our subjects were workers from six commercial colonies of *Bombus ignitus* Smith (Agrisect, Ibaraki, Japan). Colonies were maintained in nest boxes and connected to the cage through a transparent entrance tunnel and a wooden box fitted with gates, which allowed individual bees to be tested by restricting access of other bees. Inside each cage, a combination of regular fluorescent bulbs, daylight LED tubes, and UV supplemental LED bars provided illumination. The illumination intensity was maintained at 800– 1300 lux, which is above the level required for active foraging of bumble bees (ca. 700 lux, Chittka & Spaethe 2007). On non-experimental days, bees were allowed to access the cage freely between 9:00 and 20:00 for foraging; on the cage floor, we placed 2–4 plastic Petri dishes or vials, each filled with a 20% (w/w) sucrose solution and covered with a lid that had a central hole through which a cotton dental roll was inserted.

### 2.2 Artificial flowers and colour distinctions

We used artificial flowers made from clear acrylic jars (Fig. 1A; diameter = 3.5 cm, height = 4.3 cm, Muji, Tokyo). Each jar was inverted to use its base as a platform and the small depression at the center (diameter = 0.4 cm, depth = 0.2 cm) as a holder for 3 µL of 30% (w/w) sucrose solution (hereafter, nectar), which was manually added using a pipette. The surface of the jars was sanded roughly to assist bees in gripping onto the petals.

**Figure 1.**
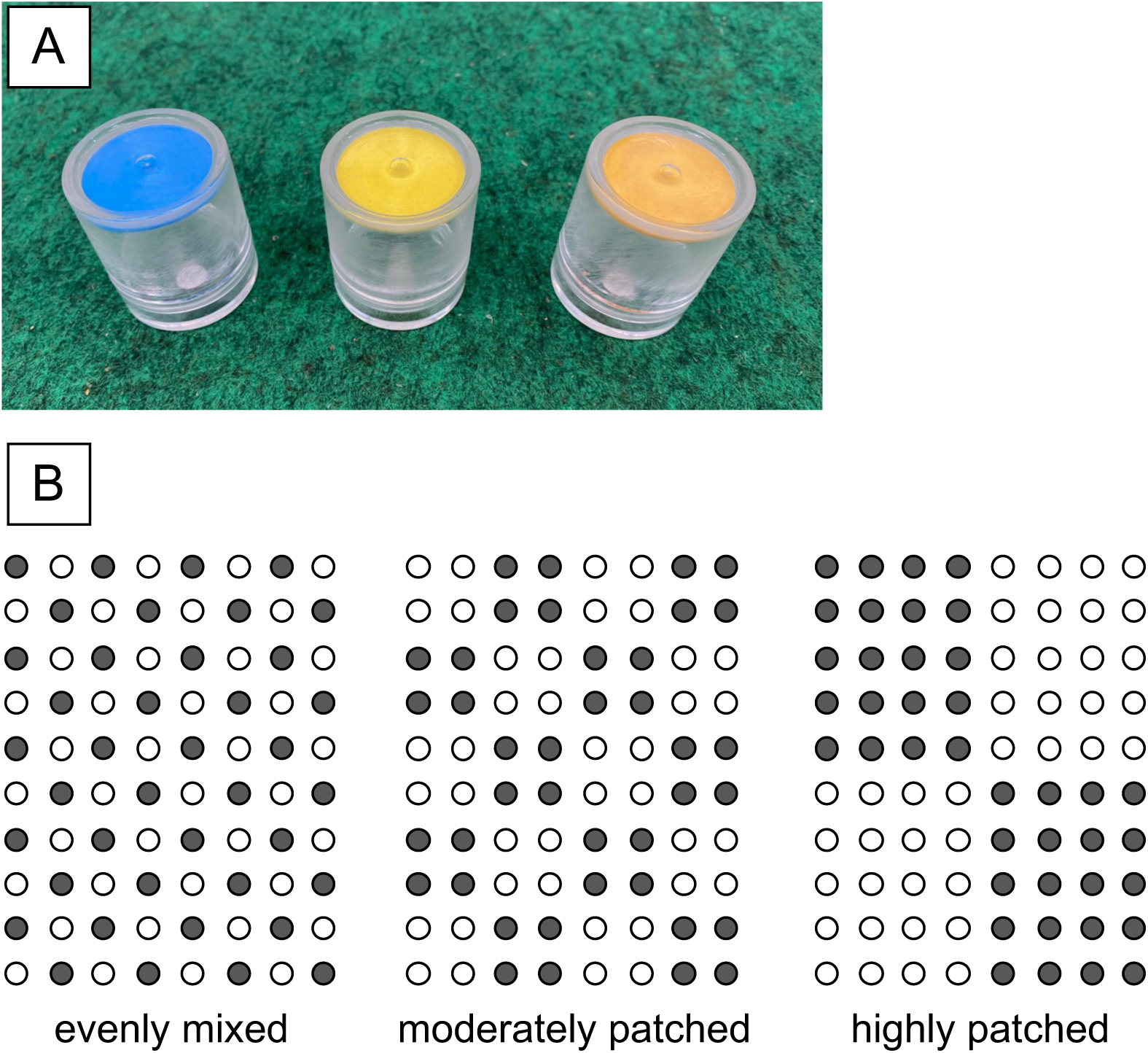
(A) Three species of artificial flowers. From left to right: blue, yellow, and golden yellow. (B) colour dimorphic arrays used in the experiment. From left to right: evenly mixed, moderately patched, and highly patched arrays. Filled and unfilled circles represent yellow and blue (or golden yellow) flowers.

We conducted two experiments to determine the effects of colour similarity on memory retrieval and realized flower constancy: the first with distinct colours, followed by the second with similar colours. For this purpose, we created three ’species’ of flowers with different colours—blue, yellow, and golden (or amber-toned) yellow —by attaching a circular piece of construction paper to the inside of each platform (Fig. 1A). We designated blue and yellow for the first experiment and yellow and golden yellow for the second experiment. These colours were selected based on perceived colour distinctions by bees. The estimations were derived from the diffuse spectral reflectance (300–700 nm) of artificial flowers and the background green carpet, measured by a spectrometer (BRC112, B&W Tek, Inc., Newar, DE, USA). The measurement was conducted relative to a white reflection standard (RS50; StellarNet, Inc., Tampa, FL, USA). A deuterium/tungsten light source (BDS100, B&W Tek, Inc.) was used for illumination. To reproduce the colours perceived by bees in our experiments, we measured the reflectance spectra of each coloured paper covered with the acrylic platform. We first translated the measured spectra into physiological inputs to the three photoreceptors of bees (UV, Blue, Green) by multiplying the irradiance spectrum of CIE standard illuminant D65 (Wyszecki & Stiles 1982) and the reported sensitivity functions of three photoreceptor classes in the retina of *Bombus terrestris* (Skorupski et al. 2007). The calculated values were standardized by those of the background green carpet, transformed into excitation values for the three receptors, and subsequently converted to x- and y-coordinates within the bee colour space, i.e., colour hexagon (Chittka 1992; Chittka & Kevan 2005). Evidence from behavioural experiments suggests that bees exhibit continuous colour perception (von Helversen 1972), with findings demonstrating that at least one species, *B. terrestris*, achieves a distinguishable accuracy rate exceeding 60% when the Euclidean distance between loci is 0.09 or greater in the hexagon unit (Dyer 2006). The colour distance between the blue and yellow flowers (0.57) is four times greater than that between the yellow and golden yellow flowers (0.15), although both pairs should be distinguishable to the bee’s eye (Fig. S1).

### 2.3 Training

Each of the two experiments comprised two consecutive days: the first day for training bees to forage from the flowers, and the second day for conducting test trials after warm-up sessions. On the first day, we arranged 40 flowers of either of the two colours, i.e., species, for the experiment in a 5 × 8 Cartesian grid on the cage floor. The distance between adjacent flowers was set to match that of test trials scheduled for the next day: 3.5 or 20 cm. We left the entrance gate open so that bees could forage freely on this array for two hours. The experimenter remained in the cage to refill the nectar using a pipette immediately after each bee visit, ensuring that a bee always encounters rewarding flowers. We subsequently switched colour to the other one of the pair and continued the training in the same manner. The order of training colours was frequently alternated between trials to prevent bias in the bees’ proximate experiences toward either of the two colours. Once bees started ’regular foraging’, wherein they would visit the flower immediately upon entering the cage, briefly return to the nest to deposit their nectar loads, and repeat the same process or foraging bout, we uniquely marked these regular foragers by gluing numbered and coloured tags onto their thoraxes. On the following day, we selected a tagged bee that was the first to approach the gate and allowed her to undergo the same training session for three foraging bouts per colour as a warm-up. After the test trial with the bee as explained below, we repeated the same warm-up sessions and subsequent tests for the other tagged bees in succession, with 1–6 bees tested per day.

### 2.4 Testing

After the warm-up session described above, we proceeded to test the bee in a mixed array of 40 flowers from each of the two species (blue and yellow for the first experiment; yellow and golden yellow for the second experiment), set up in an 8 × 10 Cartesian grid. The procedure for each test trial is identical to the preceding warm-up session, including the manual nectar refilling, with the exception of conducting only a single foraging bout on the array of 80 flowers containing equal numbers of the two species, and post-trial cleaning of the flowers with 70% ethanol to eliminate any potential effects of scent marking (Stout et al. 1998). To examine how realized flower constancy reflects potential travel costs associated with bypassing heterospecific flowers, we tested each bee with one of three arrays varying in the levels of species mixing (Fig. 1B): (1) evenly mixed, (2) moderately patched, and (3) highly patched. The relative cost of moving towards the nearest conspecific flower versus the nearest heterospecific flower, averaged across all 80 flowers, was 1.41 for the evenly mixed array, 0.98 for the moderately patched array, and 0.64 for the highly patched array, respectively. In the first test involving blue and yellow flowers, we further addressed whether the outcomes would differ if the spacing between flowers varied with the same degree of species mixing. For each array, we conducted the same tests using two distinct interflower distances: 3.5 cm and 20 cm, referred to as the high- and low-density conditions, respectively. For the second experiment, we conducted trials exclusively in the high-density condition.

In the first experiment, each bee was randomly assigned to one trial out of six conditions, which comprised combinations of three mixing levels and two density levels. In the second experiment, bees were assigned to one of three conditions representing different mixing levels. All bee behaviour during the test trials was recorded using either a video camera (HDR-SR8, SONY) or an iPhone11 (Apple), and the first 70 flower visits of each trial were used for the subsequent analyses. In total, we observed 117 bees from five colonies in the first experiment and 30 bees from one colony in the second experiment (see replication statement in Table 1).

**Table 1.** Replication statement in this study.

| Analysis | Scale of inference | Scale at which the factor of interest is applied | Number of replicates at the appropriate scale |
| --- | --- | --- | --- |
| Constancy index | Individual bees | Individual bees | First experiment (high density): 19–20 bees per mixing level<br>First experiment (low density): 19–20 bees per mixing level<br>Second experiment: 9–11 bees per mixing level |
| Number of consecutive visits to one species just before switching | Heterospecific moves | Heterospecific moves | First experiment (high density): 111–527 moves per mixing level<br>First experiment (low density): 191–1091 moves per mixing level<br>Second experiment: 111–472 moves per mixing level |
| Switch-over time | Heterospecific moves | Heterospecific moves | First experiment (high density): 100–525 moves per mixing level<br>First experiment (low density): 182–1091 moves per mixing level<br>Second experiment: 99–470 moves per mixing level |
| Decline rate in probability of switching | Interflower moves | Interflower moves | 1282 moves (pooled 19 bees at moderately patched array) |

### 2.5 Video tracking

We extracted all visit sequence and timing data from video footage. A visit to a flower was recorded when a bee landed on its platform. During the test trials, bees occasionally circled over the array or landed on the floor for a short period of time, after which they returned to the array and resumed their foraging. Each interrupted sequence was treated as two independent sets of data.

Based on the recorded visit sequences of each bee, we quantified the realized flower constancy using Gegear & Thomson’s (2004) constancy index, CI, an adaptation from Jacobs (1974), as follows:

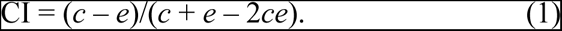

where *c* represents the observed proportion of moves made by the bee between flowers of the same species, and *e* is the proportion of moves between the same species expected based on the overall frequency of each species visited during the experiment, calculated as:

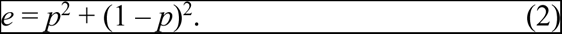

Here, *p* denotes the proportion of visits to either species of the colour pair. Thus, CI represents the degree to which an individual bee moved between flowers of the same species, while controlling for any bias in the frequency of visits to each species due to underlying colour preferences. Possible values of CI range from -1 (complete inconstancy) to +1 (complete constancy).

To test the hypothesis that the patchiness of species increases the cost of memory retrieval by facilitating consecutive visits to the same species, we also compared the switch-over time, i.e., the time taken to travel between heterospecific flowers, among arrays under different conditions. We predicted that as species patchiness increases, bees would exhibit longer consecutive visits to one species and take more time to transit between heterospecific nearest neighbours. The switch-over time was quantified as the sum of video frames (30 frames per second) required for a bee to leave one flower to alight on the next, using the open-source program Tracker version 6.1.5 (^©^2023 Douglas Brown, https://physlets.org/tracker/).

### 2.6 Statistical analysis

All analyses were performed using R version 4.1.2 (R Core Team, 2021). First, we fitted a linear mixed model (LMM) to the data to determine whether and how the level of species mixing, density, and colour contrast of paired species affected the realized flower constancy (CI). We considered species mixing level (evenly mixed/moderately patched/highly patched), density (high/low), and colour contrast (high/low) as fixed effects, with colony identity included as a random effect. Additionally, we included two interaction terms: one between mixing level and density and another between mixing level and colour contrast. Bee identity was not added as a random effect as each bee was assigned to one trial.

Next, we fitted a generalized linear mixed model (GLMM) to the data on switch-over time, employing a Poisson error distribution and a logarithmic link function. We considered species mixing level, density, and colour contrast as fixed effects, and colony and bee identity as nested random effects. Additionally, we included two interaction terms: one between mixing level and density and another between mixing level and colour contrast. Note that we exclusively used switch-over time data for cases where bees originated from a flower whose nearest neighbours included a heterospecific. This focus allowed us to eliminate possible effects of variation in the distance to the nearest heterospecifics on the switch-over time. We also asked whether species patchiness facilitates longer consecutive visits to one species before switching to another, by fitting a GLMM with a Poisson error distribution and a logarithmic link function. We considered the number of consecutive visits to one species before switching to another as response variables, species mixing level, density, and colour contrast as fixed effects, and colony and bee identity as nested random effects. Additionally, we included two interaction terms: one between mixing level and density and another between mixing level and colour contrast.

Finally, we examined whether and how the probability of switching species is affected by the accumulated number of consecutive visits to one species, aiming to understand the duration a bee could retain a search image of one species in STM when not in use. We fitted a GLMM with binomial error distribution and a logit link function to the data obtained from the moderately patched array in the first experiment, where bees were given a choice after each visit between a blue and a yellow flower, both positioned equidistant from the current flower. In this model, the response variable was dichotomous, with each movement categorized as either switching or non-switching. The number of consecutive visits to one species before each movement was considered as a fixed effect, and colony and bee identity were considered as nested random effects. Based on the coefficient and intercept estimated by GLMM, we depicted how the probability of switching species declines with the number of consecutive visits to one species.

### 2.7 Computer simulation

To determine the relative contributions of cognitive and travel costs to realized flower constancy, we developed a simulation model that generates flower visit sequences followed by a bee foraging from two species of flowers, based solely on the principle of reducing travel cost. Just like in the real experiments, we simulated bees foraging from 80 flowers, with 40 flowers of each species arranged in an 8 × 10 Cartesian grid. The two species were assumed to have identical colours perceptible to bees. This simulates a scenario in which bees incur no cognitive cost when switching species, choosing the next flower based solely on travel distances. We predicted that, in this scenario, the level of flower constancy would depend exclusively on the variation in travel cost associated with the degree of species mixing. In real bee foraging, on the other hand, bees will need to adjust their level of flower constancy to balance it with the cognitive cost, specifically the time required for memory retrieval during species switching. Therefore, by comparing the simulated results with those obtained from our experiments, we estimated the impact of memory retrieval cost on realized flower constancy.

In each simulation, a bee randomly selects a flower for its first visit. It then determines the distance it will travel based on observed probability, and subsequently selects the next flower from those located at the chosen distance. The probability of selecting a specific distance was derived from positively skewed frequency distributions of travel distances observed in warm-up sessions involving six bees (n = 1283 in high density; n = 1144 in low density; Fig. S2). During these sessions, bees foraged among an array of single-coloured flowers, providing ideal data to observe how bees travel between flowers in the absence of memory retrieval cost. Each simulation continued until the bee completed its first 70 visits, and a CI value was calculated from the generated visit sequence. In total, we generated 500 sequences (and associated CIs) for each of the six combinations of species mixing and flower density.

## 3 RESULTS

### 3.1 The first experiment using species with high colour contrast

During the experiment with blue and yellow flowers, bees consistently and markedly reduced the Constancy Index (CI) value with an increasing level of species mixing, despite the high colour contrast between the species (Fig. 2, mixing level: χ^2^ = 349.4, df = 2, P < 0.001, type-II Wald chi-squared test). As flower spacing was expanded from 3.5 to 20 cm, the CI value decreased even further, reaching levels below zero at a faster rate with increasing levels of mixing (density: χ^2^ = 75.4, df = 1, P < 0.001, mixing level × density: χ^2^ = 41.5, df = 2, P < 0.001, type-II Wald chi-squared test). The observed number of visits was 4283 of 8179 (52%) to blue flowers, and 3896 of 8179 (48%) to yellow flowers, respectively (χ^2^ = 18.3, df = 1, P < 0.001, two-tailed chi-squared test). This indicates that bees, as a group, slightly preferred blue over yellow, although the effect size was modest.

**Figure 2.**
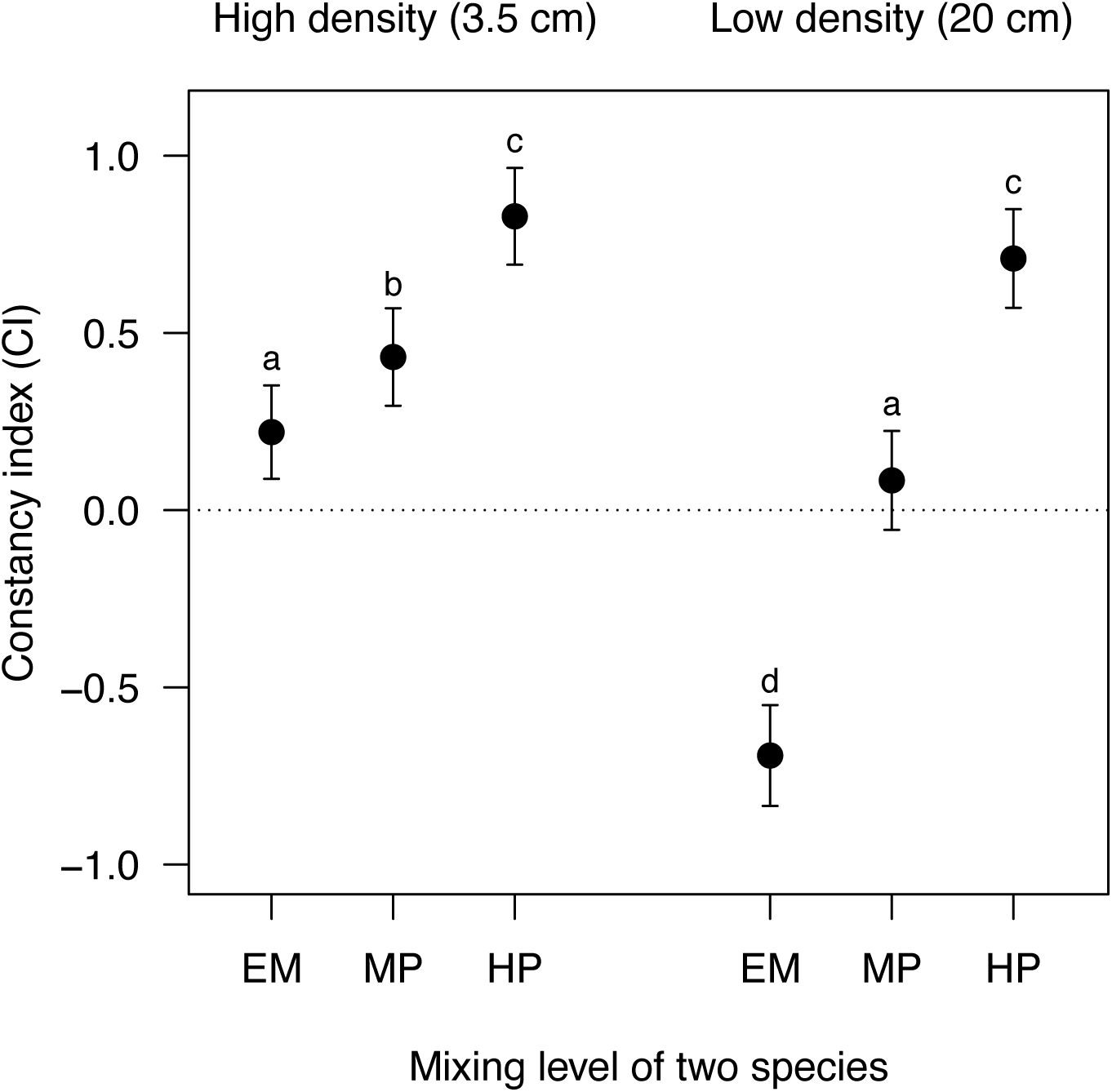
Realized flower constancy in the first experiments. Each point and error bar represent the model-adjusted mean and 95% confidential interval. EM, MP, and HP at the horizontal axis represent evenly mixed, moderately patched, and highly patched arrays, respectively. Different letters represent significant differences at the 0.05 level in pairwise tests with FDR controlling procedures (Benjamini & Hochberg 1995).

The enhanced constancy in patchy distributions further increased the cost of memory retrieval when flowers were closely spaced, potentially creating a feedback loop that amplifies the level of realized flower constancy. The number of consecutive visits to one species just before switching to another species significantly increased as species were distributed patchily, irrespective of density (Fig. 3A, mixing level: χ^2^ = 396.3, df = 2, P < 0.001, type-II Wald chi-squared test), while its overall level was significantly shorter at low density (density: χ^2^ = 47.8, df = 1, P < 0.001, type-II Wald chi-squared test). A significant interaction between density and mixing level was also detected (χ^2^ = 6.9, df = 2, P < 0.05, type-II Wald chi-squared test), although its effect sizes were negligible. In line with the tendency to make longer consecutive visits to one species, bees significantly took more time to switch species in patchy distributions (Fig. 3C, mixing level: χ^2^ = 47.6, df = 2, P < 0.001, type-II Wald chi-squared test). However, this increase in switch-over time was not observed at low density (mixing level × density: χ^2^ = 58.5, df = 2, P < 0.001, type-II Wald chi-squared test; see also the results of pairwise tests). Instead, bees consistently spent more time switching species at low density compared to high density (Fig. 3C, density: χ^2^ = 134.8, df = 1, P < 0.001, type-II Wald chi-squared test).

**Figure 3.**
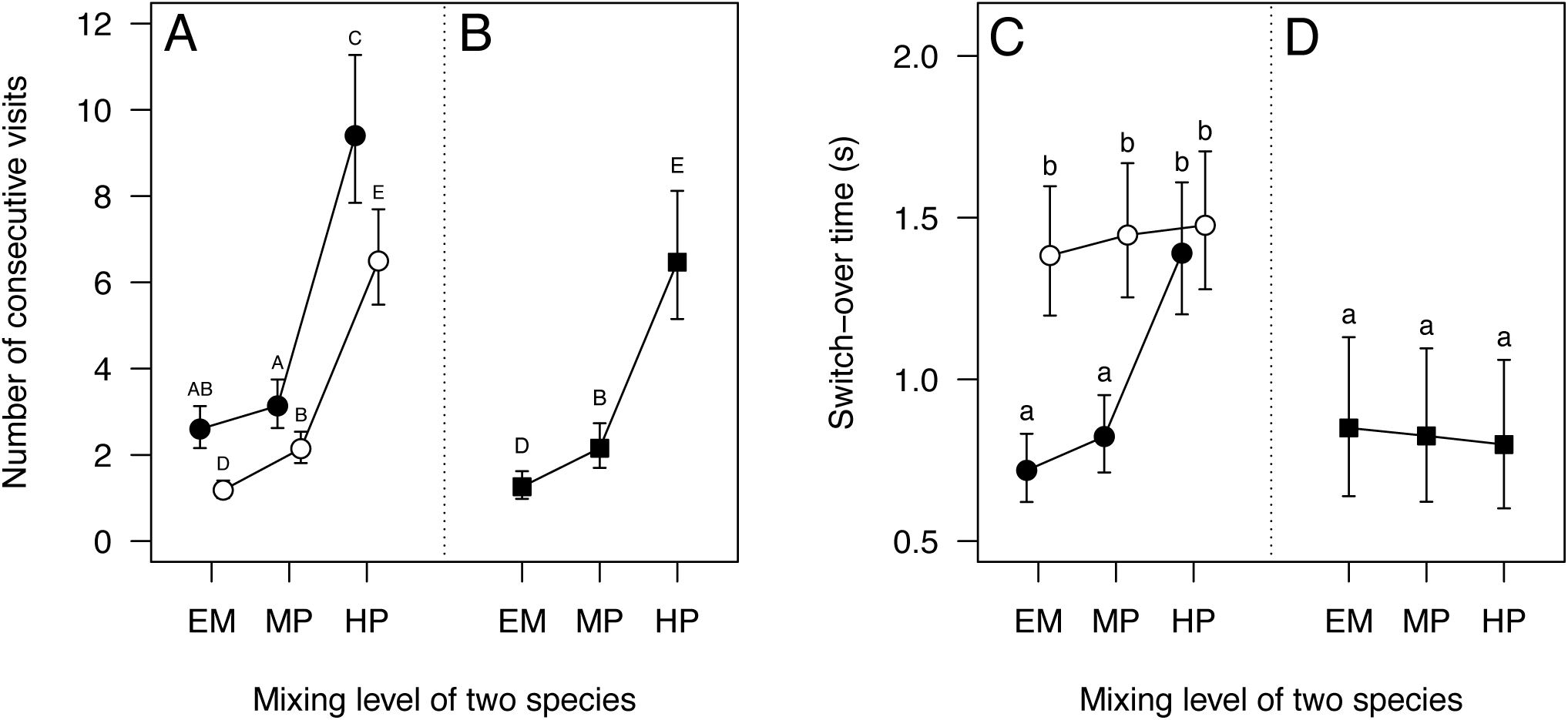
The number of consecutive visits just before switching (A, B) and switch-over time (C, D) in two experiments. The horizontal axis represents the level of species mixing. Filled and open circles represent high and low density in the first experiment, respectively. Filled squares represent the second experiment. Each symbol and error bar represents the model-adjusted mean and 95% confidential interval. Different letters represent significant differences at the 0.05 level in pairwise tests with FDR controlling procedures (Benjamini & Hochberg 1995).

Furthermore, we found more direct evidence of information decay in STM, with the probability of switching species rapidly declining as bees accumulate consecutive visits to one species (Fig. 4, χ^2^ = 25.3, df = 1, P < 0.01, type-II Wald chi-squared test). While there was a small variation among individuals—as indicated by dotted lines in Fig. 4, the overall trend suggests that the probability of switching is halved after ten consecutive visits and nearly reaches zero after 40 consecutive visits. Note that this analysis was performed solely for the high-density condition. The shorter number of consecutive visits to one species before switching observed at low density (Fig. 3A), prevented us from conducting a comparable analysis for the low-density condition.

**Figure 4.**
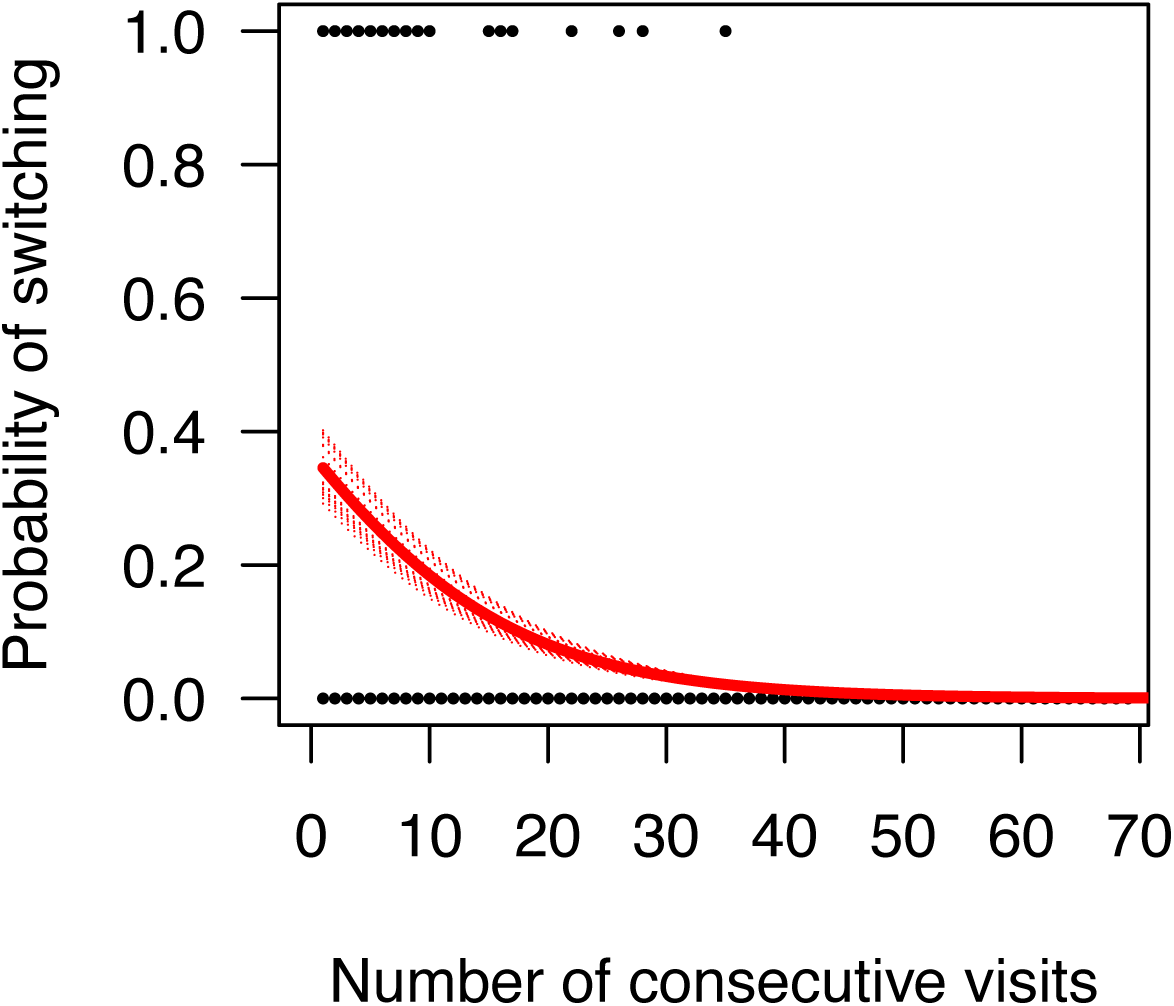
The probability of switching to another species as a function of the number of consecutive visits just before movements. Each point represents a movement (non-switching = 0, switching = 1) at high density in the first experiment. The solid line is the fitted curve from the logistic regression equation based on the coefficient and intercept estimated by GLMM: y = 1/[1 + exp(-z)], z = -0.54 + (-0.09)x. The dotted lines are fitted curves for each bee by incorporating the estimated random effect (*R_i_*) into the fitted model: y = 1/[1 + exp(-z)], z = -0.54 + *R_i_* + (-0.09)x.

### 3.2 Computer simulation and the second experiment using species with low colour contrast

The flower visit sequences generated by our computer simulations resulted in CI values that decreased sharply with increasing levels of species mixing, irrespective of density (Fig. 5). The CI values dropped below zero when species were evenly mixed, indicating that the constancy level was indistinguishable from random expectation. As depicted in Fig. 5, the simulated CI values, derived solely from bees’ preference for shorter travels, closely resembled those observed in the second experiment, where two species had similarly coloured flowers, as well as those observed at low density in the first experiment. In contrast, the observed values of CI at high density in the first experiment were quantitatively much higher than the above three sets of results. This difference was particularly pronounced for the evenly mixed array, where the CI values remained positive (Fig. 5, the results of pairwise tests). The observed number of visits was 1149 of 2072 (55%) to yellow flowers, 923 of 2072 (45%) to golden yellow flowers, respectively (χ^2^ = 24.7, df = 1, P < 0.001, two-tailed chi-squared test). This indicates that bees, as a group, slightly preferred yellow over golden yellow, although the effect size was modest.

**Figure 5.**
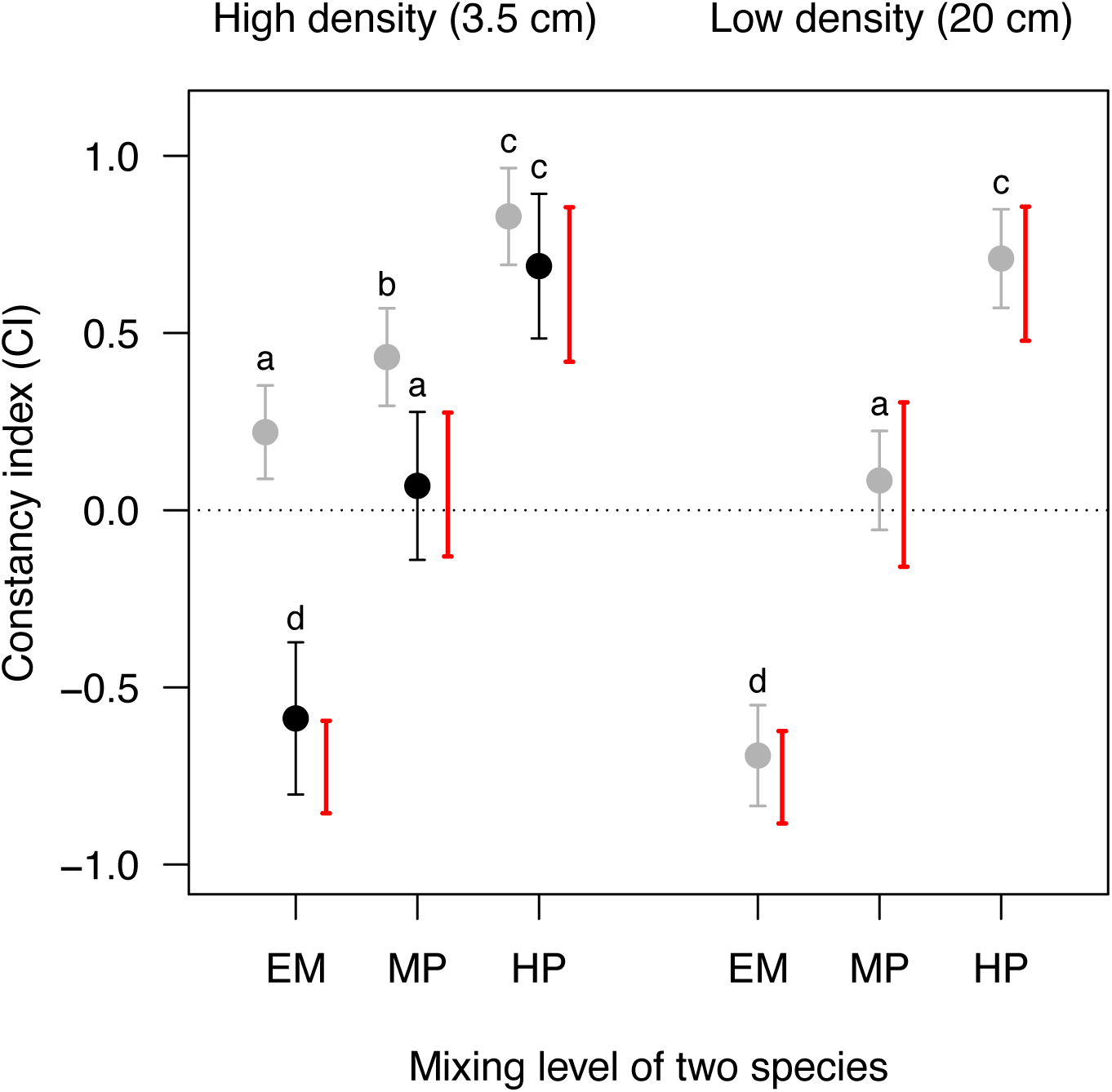
Realized flower constancy in two experiments and simulations. The horizontal axis represents the level of species mixing. Grey and black circles represent the first and second experiments, respectively. Each point and error bar represent the model-adjusted mean and 95% confidential interval. Different letters represent significant differences at the 0.05 level in pairwise tests with FDR controlling procedures (Benjamini & Hochberg 1995). Each red bar represents the 5th to 95th percentile range of 500 CI values generated by the simulation.

As for the number of consecutive visits to one species just before switching, the results of the second experiment were statistically indistinguishable from those at low density in the first experiment (Fig. 3A and B, the results of pairwise tests). Despite this increase in consecutive visits, switch-over time did not increase with the levels of species mixing (Fig. 3D, mixing level × colour contrast: χ^2^ = 55.7, df = 2, P < 0.001, type-II Wald chi-squared test; see also the results of pairwise tests). Instead, bees consistently switch species as quickly as those in the evenly mixed array at high density (Fig. 3C and D, the results of pairwise test).

## 4 DISCUSSION

To the best of our knowledge, this study is the first to provide compelling evidence that realized flower constancy in pollinators is not merely a result of cognitive limitation, but rather an adaptive behaviour that strategically balances cognitive and travel costs across different environmental contexts. Below, we will first discuss our key findings in the two experiments. Building upon these results, we will develop a conceptual framework for understanding how realized flower constancy varies across environments, drawing upon principles from optimal foraging theory. In line with this framework, we will finally propose a novel perspective on conditions under which diverse floral phenotypes benefit plants.

### 4.1 Realized flower constancy as an integrated response to species mixing, spacing, and floral colour difference

Throughout the first experiment, bees dramatically decreased realized flower constancy as species mixing increased (Fig. 2). This aligns with our prediction; however, considering that the distinct colours of blue and yellow flowers have been thought to impose the highest cost of memory retrieval (Chittka et al. 2001), such a substantial reduction in flower constancy exceeds our expectation. Since our arrays vary only in the positional relationship of the two species, while nearest-neighbour distance remains constant, we can attribute the decrease in realized flower constancy to bees’ preference for short distances, i.e., their aversion to travel cost (Pyke 1978). In other words, bees would have engaged in more frequent species switching at the cost of memory retrieval in order to mitigate excessive travel costs. As far as we know, one study in the literature has attempted to examine the effects of travel cost as a potential counteracting force against the cost of memory retrieval (Gegear & Thomson 2004). Unfortunately, these authors chose a dimorphic array for their experiments, specifically designed to detect only the effects of memory retrieval cost by providing bees with an equal choice of two flower species (both nearest and second-nearest neighbours) upon leaving any flower. With this setup, the effects of variation they introduced for inter-flower distance on flower constancy would be minimal, as travel costs do not vary with the bees’ foraging selectivity.

When flowers were spaced farther apart, realized flower constancy dropped to levels indistinguishable from those predicted by the simulation model under the assumption of no memory retrieval cost (Figs. 2 & 5). Given that high- and low-density arrays did not differ in the positional relationships of the two species, the observed difference in the constancy levels cannot be attributed to either memory retrieval or travel costs. One probable reason for the reduced constancy with increased spacing may be the limited ability of bumble bees to detect a specific size of visual target from a distance (Ishii & Masuda 2014). When flowers are more spaced out, bees are less likely to detect them until they get closer. This may apply to our low-density condition, where the minimum target size that bees could detect from a distance of 20 cm is 1.9 cm (calculated from the minimum visual angle of 5–6° reported for *Bombus terrestris* in Spaethe & Chittka 2003). Considering that our flower is coloured only on its platform, it is highly likely that its apparent size, seen from an angle, is smaller than 1.9 cm. In this condition, bees will need extra time to find any of the surrounding flowers. As a result, information on the last visited flower in a bee’s STM would fade back into LTM before the bee finds other flowers. Indeed, visual stimuli retrieved from LTM to STM have been indicated to last only for 1–2 s in bumble bees (Raine & Chittka 2007). Such decay of information after each visit would minimize the difference in memory retrieval cost between the two species and increase the likelihood of species switching.

While memory decay from STM to LTM at low density leads to a decrease in realized flower constancy, the same memory attenuation may strengthen constancy levels at high density due to bees’ tendency to move between neighbouring conspecifics. This could have occurred in the highly patched distribution where bees made longer consecutive visits to one species as a direct consequence of bees’ preference to travel short distances (Fig. 3A). During these persistent visits to a single species, bees would not have been able to maintain the memory of another species in STM and prevent it from decaying into LTM, leading to an escalation in retrieval cost. Our observations that bees took more time to switch species in the highly patched array at high density (Fig. 3C), as well as being less inclined to switch species after accumulating consecutive visits to one species (Fig. 4), support this conjecture. Such a feedback effect of constant visits on the cognitive cost of switching species may explain why bees foraging in monospecific meadows were more tenacious to the last visited species, even when offered another species (Wilson & Stine 1996). Note, however, that the increased cost of memory retrieval in patchily distributed flowers will be canceled out at low density, where the memory of both species fades back into LTM while bees are searching to meet the visual detection limit (see above discussion). To support this view, the switch-over time did not vary significantly across different levels of species mixing at low density (Fig. 3C).

One point that might be worth discussing is the reason why the decline in the probability of switching was gradual rather than an abrupt drop (Fig. 4), given that information about the last visited flower fades from STM to LTM within a few seconds. This may be explained by the multilayered nature of bees’ memory system. While we followed the traditional dichotomy between STM and LTM for the sake of simplicity (Chittka et al. 1999), bees in the real world have been suggested to possess multiple phases within the two components of memory, which differ from one another in their durations and capacities (Menzel 1999). Hence, the gradual decline of the probability of switching may reflect that information about the last visited species transferred from STM to longer-term phases of memory incrementally, rather than solely indicating the duration for which the information is retained in STM.

When bees encountered two species with similar floral colours in the second experiment, the levels of realized flower constancy dropped significantly more, reaching levels indistinguishable from those obtained from the simulation assuming cognitive costs (Fig. 5). This strongly suggests that bees experienced minimal or no memory retrieval costs when switching between similar colours and their decisions regarding the next flower were primarily aimed at minimizing travel costs. The bees likely perceived similarly coloured flowers as belonging to the same species, thereby operating with a shared search image, without needing to retrieve distinct information from LTM to STM each time they switched species. It is also possible that they learned to generalize between the two equally rewarding colour stimuli (Gumbert 2000; Benard et al. 2006), even though they are capable of distinguishing them, thereby successfully avoiding unnecessary cognitive load. In line with this interpretation, the switch-over time in the second experiment remained consistently low irrespective of the mixing levels (Fig. 3D), whereas the number of consecutive visits to one species increased with its patchiness (Fig. 3B). Chittka et al. (2001) also reported that flower constancy exhibited by five species of apid bees increased with bee-subjective colour distance following a non-linear curve, with a sharp rise occurring at a narrow range of colour distance. These observations suggest that alternating two species of flowers exerts cognitive costs of switching memories only when their phenotypic difference exceeds the level below which bees can manage them with a shared memory.

### 4.2 Conceptual framework for understanding realized flower constancy

The findings presented here collectively support our initial hypothesis: realized flower constancy in bees is not solely determined by cognitive limitation imposed by interspecific differences in floral phenotypes. Instead, it reflects an optimal foraging strategy that dynamically varies across environments, balancing cognitive and travel costs. Based on these findings, here we propose a conceptual framework for understanding how pollinators determine the level of realized flower constancy in a certain environment.

The four diagrams in Fig. 6 are graphical representations illustrating how cognitive and travel costs for pollinators would vary while foraging in a mixture of species, based on the level of flower constancy. They also depict how these relationships would change with the degrees of phenotypic differences and spatial mixing between plant species. Note that only the relationship between travel cost and constancy level changes with the degree of species mixing (columns), while only the relationship between memory retrieval cost and constancy level changes with phenotypic differences (rows). The triangle in each condition represents realized flower constancy an optimal forager would choose to balance cognitive and travel costs and minimize their total.

**Figure 6.**
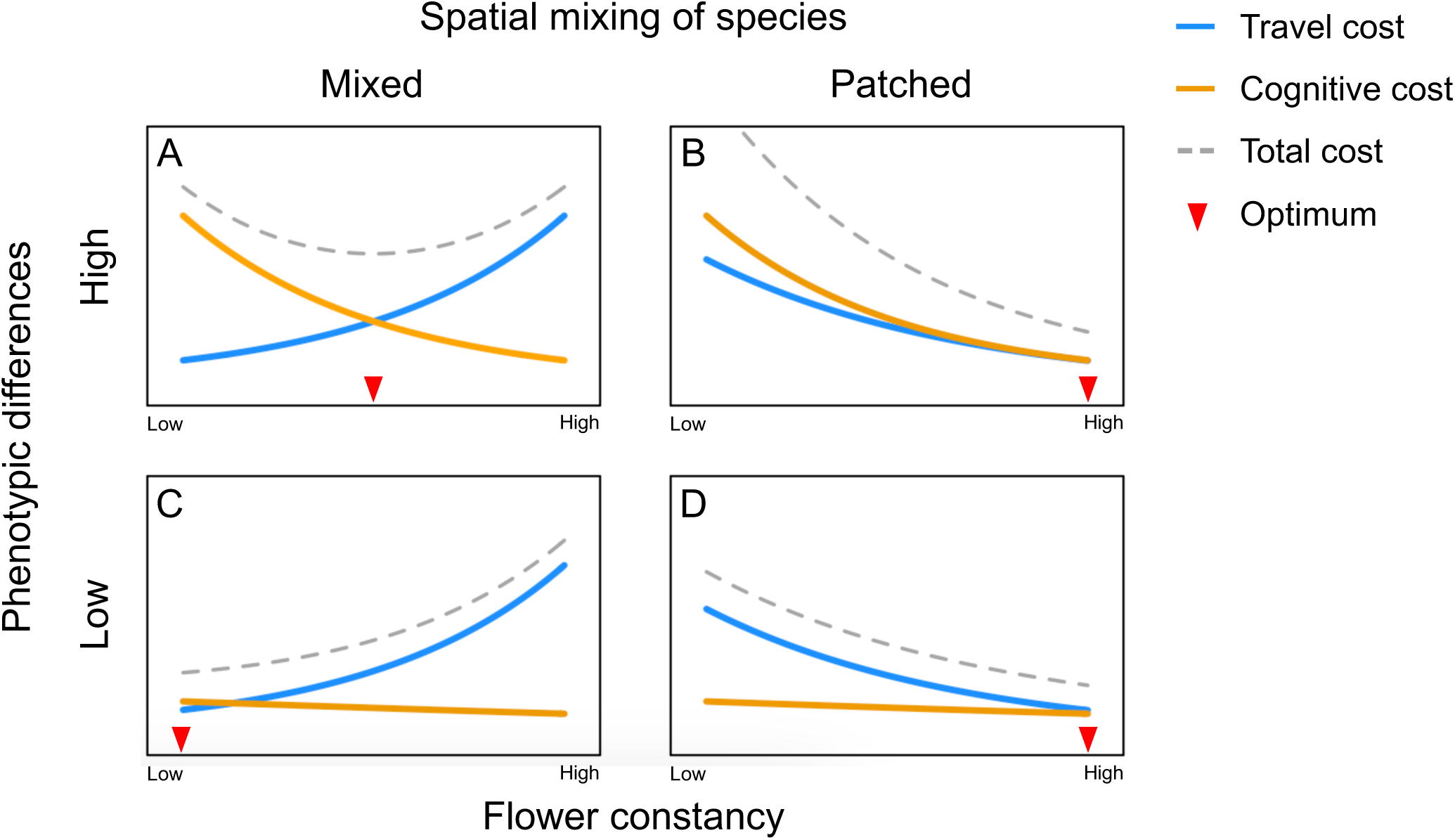
The conceptual framework for comprehending the variation in realized flower constancy across environments characterized by different combinations of phenotypic differences (rows) and spatial species mixing (columns). Blue, solid orange solid, and grey dotted curves represent the changes in travel cost, cognitive cost, and total cost respectively, corresponding to varying levels of flower constancy. The red inverted triangles depict the optimal (i.e., realized) levels of flower constancy that minimize the total cost, combining time for memory retrieval and travel. Note that phenotypic differences significantly affect realized flower constancy only when plant species are spatially mixed (A and C). In contrast, phenotypic differences have a minimum impact on realized flower constancy when species are patchily distributed (B and D).

When different plant species are highly mixed in space (A, C), more constant foragers benefit from the reduced cost of memory retrieval (orange curves), while incurring a significantly increased travel cost due to more frequent skips of heterospecific neighbours (blue curves). When two species have distinct floral phenotypes, the optimal level of realized flower constancy will be intermediate, so as to balance between cognitive and travel costs (A). When floral phenotypes are similar between species, pollinators will incur minimal or no cognitive costs. In this situation, optimal foragers should exhibit the lowest levels of realized flower constancy (C). On the contrary, when plant species are patchily distributed (B, D), more constant foragers incur a significantly lower travel cost due to the reduced skips of heterospecific neighbours, while maintaining the benefit of avoiding the cost of memory retrieval. In this condition, the optimal level of realized flower constancy should be highest, irrespective of the degree of interspecific phenotypic differences, because selective foraging could minimize both cognitive and travel costs without any conflict. It should be noted that graphs C and D also correspond to situations where plant species are spaced out, resulting in a leveling off of the difference in the cost of memory retrieval (Fig. 3C). These considerations align closely with the observed changes in realized flower constancy across species mixing and colour similarity in our experiments (Fig. 5). The proposed framework will help predict the levels of realized flower constancy by pollinators in natural environments where phenotypic differences and spatial relationships among plant species vary substantially.

### 4.3 When do diverse floral phenotypes benefit plants?

Our conceptual framework may explain the lack of consistency in the literature regarding floral colour dissimilarity among co-flowering species (Gumbert et al. 1999; de Jager et al. 2011; Makino & Yokoyama 2015; Shrestha et al. 2019). Previous authors have implicitly or explicitly assumed that the increased phenotypic dissimilarity among co-flowering species would discourage pollinators from moving between these species. Our findings, however, clearly demonstrate that this is not always the case when considering the costs of pollinator movement due to the positional relationships among plant species.

Notably, as observed in bumblebees, the realized flower constancy exhibited by pollinators is expected to increase with differences in floral phenotype only when multiple plant species are spatially intermingled. When species are patchily distributed, pollinators would become more constant to one species, irrespective of how species are phenotypically different from one another. In other words, spatial mixing of multiple plant species may enhance the benefits of having distinct floral phenotypes from co-flowering species by reducing interspecies pollinator movements, thereby promoting floral diversity within a single plant community through ecological and evolutionary processes.

In conclusion, the findings from our indoor experiments with bumble bees shed new light on the dynamics of realized flower constancy in pollinating insects. Contrary to previous assumptions, we discovered that realized flower constancy is not solely a result of cognitive limitation but rather a flexible foraging strategy that optimizes the balance between cognitive and travel costs. Our results demonstrate that realized flower constancy (1) dramatically decreases as the spatial mixing of plant species increases, (2) increases due to a rapid decay of information about one species of flower from short-term memory to long-term storage while pollinators are visiting the other species consecutively, and (3) is reduced to a minimum level when plant species with similarly coloured flowers are evenly mixed, while it is hardly affected by colour similarity when species are patchily distributed. These results highlight the importance of spatial mixing of species in understanding the evolution and maintenance of floral diversity within plant communities.

## Acknowledgments

We are grateful to the members of the Ecological Interactions Laboratory at the University of Tsukuba for their invariable support throughout this study; the staff of the Botanical Garden at the University of Tsukuba for allowing us to use their laboratory space; and Yukie Sato and Kazuichi Sakamoto for insightful discussions. This work was supported by a JSPS Grants-in-Aid for Scientific Research (KAKENHI no. 19K06834) and a field research project of the Mountain Science Center at the University of Tsukuba.

## References

Benard, J., Stach, S., & Giurfa, M. (2006). Categorization of visual stimuli in the honeybee *Apis mellifera*. Animal Cognition, 9, 257–270. 10.1007/s10071-006-0032-9

Benjamini, Y., & Hochberg, Y. (1995). Controlling the false discovery rate: a practical and powerful approach to multiple testing. Journal of the Royal Statistical Society B, 57, 289–300. 10.1111/j.2517-6161.1995.tb02031.x

Chittka, L. (1992). The colour hexagon: a chromaticity diagram based on photoreceptor excitations as a generalized representation of colour opponency. Journal of Comparative Physiology A, 170, 533–543. 10.1007/BF00199331

Chittka, L., Gumbert, A., & Kunze, J. (1997). Foraging dynamics of bumble bees: correlates of movements within and between plant species. Behavioral Ecology, 8, 239–249. 10.1093/beheco/8.3.239

Chittka, L., & Kevan, R. G. (2005). Flower colour as advertisement. In Dafni, A., Kevan, P. G., & Husband. B. C. (Eds.), Practical Pollination Biology (pp. 157–196). Environquest Ltd., Cambridge, Ontario.

Chittka, L., & Spaethe, J. (2007). Visual search and the importance of time in complex decision making by bees. Arthropod-Plant Interactions, 1, 37–44. 10.1073/pnas.0501440102

Chittka, L., Spaethe, J., Schmidt, A., & Hickelsberger, A. (2001). Adaptation, constraint, and chance in the evolution of flower color and pollinator color vision. In Chittka, L., & Thomson, J. D. (Eds), Cognitive Ecology of Pollination (pp. 106–126). Cambridge University Press, Cambridge.

Chittka, L., Thomson, J. D., & Waser, N. M. (1999). Flower constancy, insect psychology, and plant evolution. Naturwissenschaften, 86, 361–377. 10.1007/s001140050636

De Jager, M. L., Dreyer, L. L., & Ellis, A. G. (2011). Do pollinators influence the assembly of flower colours within plant communities? Oecologia, 166, 543–553. 10.1007/s00442-010-1879-7

Dyer, A. G. (2006). Bee discrimination of flower colours in natural settings by the bumblebee species *Bombus terrestris* (Hymenoptera: Apidae). Entomologies Generalis, 28, 257–268. 10.1127/entom.gen/28/2006/257

Gegear, R. J., & Laverty, T. M. (2005). Flower constancy in bumblebees: A test of the trait variability hypothesis. Animal Behaviour, 69, 939–949. 10.1016/j.anbehav.2004.06.029

Gegear, R. J., & Thomson, J. D. (2004). Does the flower constancy of bumble bees reflect foraging economics? Ethology, 110, 793–805. 10.1111/j.1439-0310.2004.01010.x

Goulson, D. (2000). Are insects flower constant because they use search images to find flowers? Oikos, 88, 547–552. 10.1034/j.1600-0706.2000.880311.x

Greggers, U., & Menzel, R. (1993). Memory dynamics and foraging strategies of honeybees. Behavioral Ecology and Sociobiology, 32, 17–29. 10.1007/BF00172219

Grüter, C., & Ratnieks, F. L. (2011). Flower constancy in insect pollinators: adaptive foraging behaviour or cognitive limitation? Communicative & Integrative Biology, 4, 633–636. 10.4161/cib.16972

Gumbert, A., Kunze, J., & Chittka, L. (1999). Floral colour diversity in plant communities, bee colour space and a null model. Proceedings of the Royal Society B, 266, 1711–1716. 10.1098/rspb.1999.0836

Gumbert, A. (2000). Color choices by bumble bees (*Bombus terrestris*): innate preferences and generalization after learning. Behavioral Ecology and Sociobiology, 48, 36–43. 10.1007/s002650000213

Hopkins, R., & Rausher, M. D. (2012). Pollinator-mediated selection on flower color allele drives reinforcement. Science, 335, 1090–1092. 10.1126/science.1215198

Ishii, H. S. (2005). Analysis of bumblebee visitation sequences within single bouts: Implication of the overstrike effect on short-term memory. Behavioral Ecology and Sociobiology, 57, 599–610. 10.1007/s00265-004-0889-z

Ishii, H. S. (2006). Floral display size influences subsequent plant choice by bumble bees. Functional Ecology, 20, 233–238. 10.1111/j.1365-2435.2006.01106.x

Ishii, H. S., & Kadoya, E. Z. (2016). Legitimate visitors and nectar robbers on *Trifolium pratense* showed contrasting flower fidelity versus co-flowering plant species: could motor learning be a major determinant of flower constancy by bumble bees? Behavioral Ecology and Sociobiology, 70, 377–386. 10.1007/s00265-016-2057-7

Ishii, H. S., & Masuda, H. (2014). Effect of flower visual angle on flower constancy: a test of the search image hypothesis. Behavioral Ecology, 25, 933–944. 10.1093/beheco/aru071

Jacobs, J. (1974). Quantitative measurement of food selection: a modification of the forage ratio and Ivlev’s electivity index. Oecologia, 14, 413–417. 10.1007/BF00384581

Jones, K. N. (2001). Pollinator-mediated assortative mating: causes and consequences. In Chittka, L., & Thomson, J. D. (Eds), Cognitive Ecology of Pollination (pp. 259–273). Cambridge University Press, Cambridge.

Lewis, C. A. (1986). Memory constraints and flower choice in *Pieris rapae*. Science, 232, 863–865. 10.1126/science.232.4752.863

Makino, T. T., & Yokoyama, J. (2015). Nonrandom composition of flower colors in a plant community: mutually different co-flowering natives and disturbance by aliens. PLoS ONE, 10(12), e0143443. 10.1371/journal.pone.0143443

Menzel, R. (1999). Memory dynamics in the honeybee. Journal of Comparative Physiology A, 323–340. 10.1007/s003590050392

Morales, C. L., & Traveset, A. (2008). Interspecific pollen transfer: magnitude, prevalence and consequences for plant fitness. Critical Review in Plant sciences, 27, 221–238. 10.1080/07352680802205631

Moreira-Hernández, I. J., & Muchhala, N. (2019). Importance of pollinator-mediated interspecific pollen transfer for angiosperm evolution. Annual Review of Ecology, Evolution, and Systematics, 50, 191–217. 10.1146/annurev-ecolsys-110218-024804

Ohashi, K., Thomson, J. D., & D’Souza, D. (2007). Trapline foraging by bumble bees: IV. Optimization of route geometry in the absence of competition. Behavioral Ecology, 18, 1–11. 10.1093/beheco/arl053

Pyke, G. H. (1978). Optimal foraging: movement patterns of bumblebees between inflorescences. Theoretical Population Biology, 13, 72–98. 10.1016/0040-5809(78)90036-9

Pyke, G. H. (1984). Optimal foraging theory: A critical review. *Annual Review of Ecology*, Evolution and Systematics, 15, 523–575. 10.1146/annurev.es.15.110184.002515

R Core Team. (2021). R: A language and environment for statistical computing. R Foundation for Statistical Computing. https://www.R-project.org

Raine, N. E., & Chittka, L. (2007). Flower constancy and memory dynamics in bumblebees (Hymenoptera: Apidae: *Bombus*). Entomologies Generalis, 29, 179–199. 10.1127/entom.gen/29/2007/179

Shrestha, M., Dyer, A. G., Garcia, J. E., & Burd, M. (2019). Floral colour structure in two Australian herbaceous communities: It depends on who is looking. Annals of Botany, 124, 221–232. 10.1093/aob/mcz043

Skorupski, P., Doering, T., & Chittka, L. (2007). Photoreceptor spectral sensitivity in island and mainland populations of bumblebee, Bombus terrestris. Jouranl of Comparative Physiology A, 193, 485–494. 10.1007/s00359-006-0206-6

Spaethe, J., & Chittka, L. (2003). Interindividual variation of eye optics and single object resolution in bumblebees. Journal of Experimental Biology, 206, 3447–3453. 10.1242/jeb.00570

Stout, J. C, Goulson, D., & Allen, J. A. (1998). Repellent scent-marking of flowers by a guild of foraging bumblebees (*Bombus* spp.). Behavioral Ecology and Sociobiology, 43, 317–326. 10.1007/s002650050497

von Helversen, O. (1972). The relationship between difference in stimuli and choice frequency in training experiments with the honey bee. In Wehner, R. (Eds), Information processing in the visual system of arthropods (pp. 323–334). Springer, Berlin.

Wilson, P., & Stine, M. (1996). Floral constancy in bumble bees: handling efficiency or perceptual conditioning? Oecologia,106, 493–499. 10.1007/BF00329707

Wyszecki, G., & Stiles, W. S. (1982). Color science, concepts and methods, quantitative data and formulae, 2nd edition. New York.

