## Supporting information for "Realized flower constancy in bumble bees: optimal foraging strategy balancing cognitive and travel costs and its possible consequences for floral diversity"

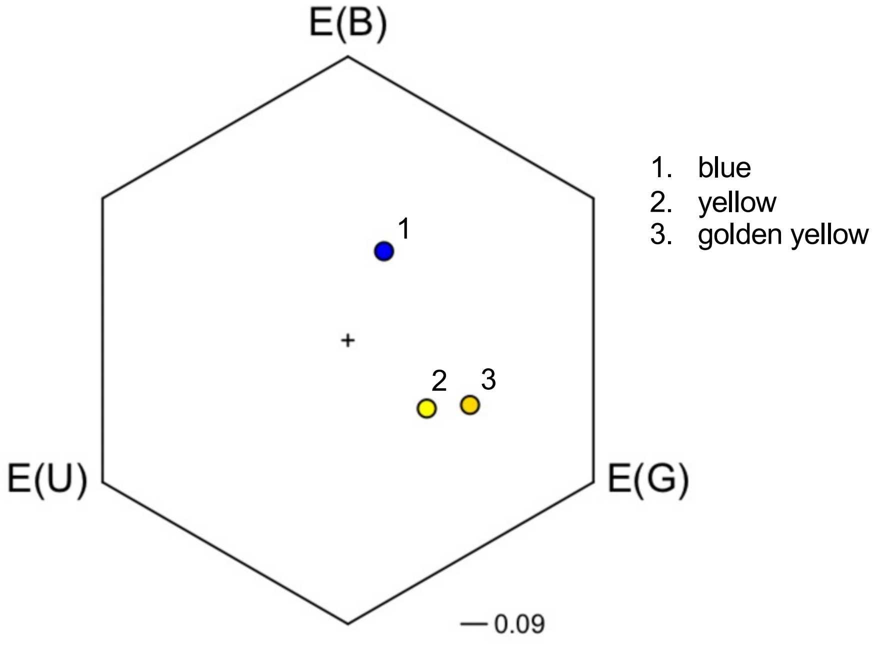


**Figure S1** Colour loci of artificial flowers in the bee colour space, i.e., colour hexagon (Chittka 1992). Three of the axes correspond to excitation values of photoreceptors sensitive to ultraviolet E(U), blue E(B), and green E(G). The angular position from the centre (the loci of the green carpet background) corresponds to hue perceived by bees. The distance between the centre and any vertex is 1 and colours that differ by distances above 0.09 are distinguishable for bees (Dyer 2006).

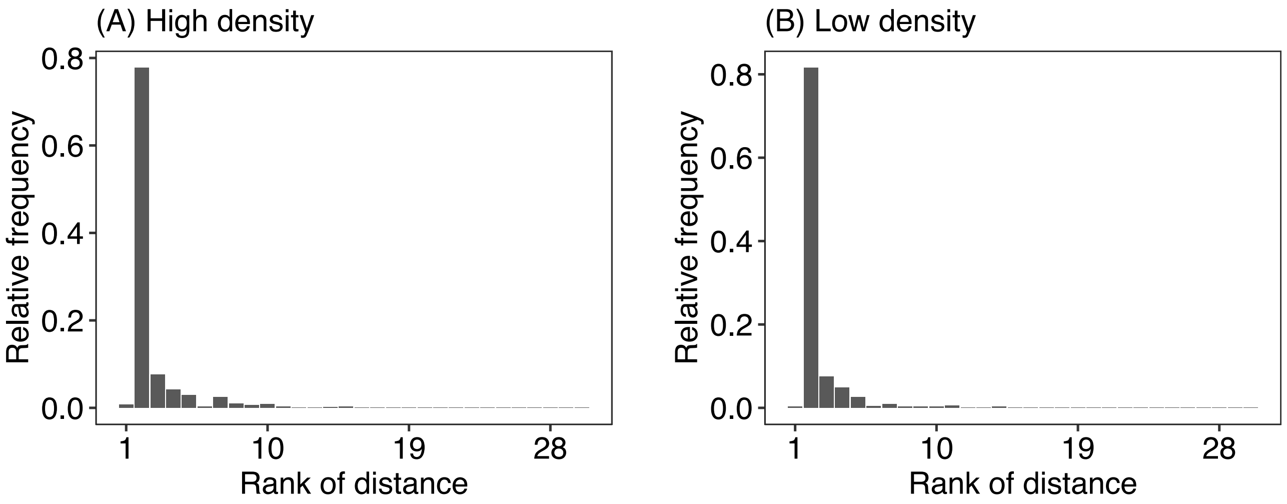


**Figure S2** Relative frequency distributions of travel distance derived from the warm-up sessions at (A) high- and (B) low-density conditions. The horizontal axis represents the rank of travel distance. Note that rank 2 refers to the case in which bees selected the nearest flower. Rank 1 refers to the case in which bees revisited the flower last visited. In simulations, the relative frequency value for each rank was used as the probability that bees select a specific distance.

**References**

Chittka, L. (1992). The colour hexagon: a chromaticity diagram based on photoreceptor excitations as a generalized representation of colour opponency. *Journal of Comparative Physiology A*, 170, 533–543. https://doi.org/10.1007/BF00199331

Dyer, A. G. (2006). Bee discrimination of flower colours in natural settings by the bumblebee species *Bombus terrestris* (Hymenoptera: Apidae). *Entomologies Generalis*, 28, 257–268. https://doi.org/10.1127/entom.gen/28/2006/257
